# Rubsicolins are naturally occurring G-protein-biased delta opioid receptor peptides

**DOI:** 10.1101/433805

**Authors:** Robert J. Cassell, Kendall L. Mores, Breanna L. Zerfas, Amr H. Mahmoud, Markus A. Lill, Darci J. Trader, Richard M. van Rijn

## Abstract

The impact that β-arrestin proteins have on G-protein-coupled receptor trafficking, signaling and physiological behavior has gained much appreciation over the past decade. A number of studies have attributed the side effects associated with the use of naturally occurring and synthetic opioids, such as respiratory depression and constipation, to excessive recruitment of β-arrestin. These findings have led to the development of biased opioid small molecule agonists that do not recruit β-arrestin, activating only the canonical G-protein pathway. Similar G-protein biased small molecule opioids have been found to occur in nature, particularly within kratom, and opioids within salvia have served as a template for the synthesis of other G-protein-biased opioids. Here, we present the first report of naturally occurring peptides that selectively activate G-protein signaling pathways with minimal β-arrestin recruitment. We find that rubiscolin peptides, which are produced as cleavage products of the plant protein rubisco, bind to and activate G-protein signaling at δ opioid receptors. However, unlike the naturally occurring δ opioid peptides leu-enkephalin and deltorphin II, the rubiscolin peptides only very weakly recruit β-arrestin 2 and have undectable recruitment of β-arrestin 1 at the δ opioid receptor.

## Introduction

Opioid receptors are G protein-coupled receptors (GPCRs) that are widely expressed throughout the body and regulate a diverse array of physiological functions, including pain sensation, respiration, mood and reward. Opioid receptors are activated by endogenous opioids, such as endorphins and enkephalins, by natural products such as *Papaver somniferum* (opium) and *Salvia divinorum*, and by synthetic opioids such as fentanyl and methadone. The current clinically-used opioids provide strong analgesic support in operative and post-operative settings, but their use is also associated with adverse effects including the development of tolerance, constipation, opioid use disorder and respiratory depression, which can ultimately prove fatal.

Studies utilizing β-arrestin 2 knockout mice revealed that certain opioid side effects may be associated with β-arrestin 2 recruitment upon μ-opioid receptor activation (Raehal et al., 2005). These early studies ignited efforts to discover or develop opioids which exclusively activate G-protein signaling without also eliciting β-arrestin recruitment. Such G-protein biased small molecules, like TRV130 (DeWire et al., 2013) and PZM21 (Manglik et al., 2016), have recently been synthesized and indeed appear to possess a reduced side effect profile. G-protein biased opioids also appear to occur naturally, as revealed by the pharmacological characterization of opioids found within *Mitragynina speciosa* (kratom) (Varadi et al., 2016). Additionally, naturally occurring opioids can serve as the basis of G-protein-biased semi-synthetic opioids, as is the case for herkinorin which is derived from naturally occurring salvinorin (Groer et al., 2007).

In 2001 peptides were identified from spinach rubisco (ribulose-1,5-bisphosphate carboxylase/oxygenase), a 557 kDa enzyme highly common in photosynthetic organisms, and it was found that two rubisco peptides YPLDL (rubiscolin-5) and YPLDLF (rubsicolin-6) have micromolar affinity and potency at δ opioid receptors (δ-ORs) *in vitro*. Interestingly a study investigating the effect of rubiscolin on skin inflammation also found that rubiscolin-6 prevented δ-OR internalization (Chajra et al., 2015). Internalization of δ-ORs tends to be correlated with β-arrestin recruitment (Pradhan et al., 2009), and thus the aforementioned reduction of δ-OR internalization with rubiscolin-6 may suggest that rubiscolin peptides do not recruit β-arrestin to δ-ORs. This led us to hypothesize that these rubiscolin peptides possess G-protein-bias when compared with the enkephalin and deltorphin-classes of peptides, which are known to recruit β-arrestin (Chiang et al., 2016; Molinari et al., 2010). Using cellular signaling assays we identified that the rubiscolin peptides indeed have a lower propensity to recruit β-arrestin than leu-enkephalin and deltorphin II, while acting as full agonists in a G-protein-mediated cAMP assay.

## Methods

### Drugs and chemicals

[D-Ala^2^]-deltorphin II was purchased from Tocris, R&D systems (Minneapolis, MN, USA). Naltrindole hydrochloride, forskolin, leu-enkephalin, and piperidine were purchased from Sigma-Aldrich (St. Louis, MO, USA). [^3^H]DPDPE (lot 2399128, 49.8 Ci/mml) was purchased from Perkin Elmer (Waltham, MA). Fmoc-protected amino acids were purchased from NovaBiochem (Billerica, MA), Fmoc-L-Leu-Wang resin (100-200 mesh, 1% divinylbenzene, 0.745 meq/g) and Fmoc-L-Phe-Wang resin (200-400 mesh, 1% divinylbenzene, 0.443 meq/g) were purchased from Chem-Impex International (Wood Dale, IL). (1H-Benzotriazol-1-yloxy)(dimethylamino)-N,N-dimethylmethaniminium hexafluorophosphate (HBTU) was purchased from Oakwood Chemical (N. Estill, SC), N,N-dimethylformamide, dichloromethane, N,N-diisopropylethylamine and acetonitrile (HPLC-grade) were purchased from Fisher Scientific (Hampton, NH), trifluoroacetic acid was purchased from VWR Chemicals (Radnor, PA) and triisopropylsilane was purchased from Alfa Aesar (Ward Hill, MA). All reagents were used as received with no further purification. For the cellular assays, all peptides were dissolved in sterile water.

### Synthesis and characterization of rubiscolin peptides

Synthesis of both peptides was performed on a 0.022 mmol scale. Peptides were synthesized using standard Fmoc-based solid-phase synthesis with HBTU. Fmoc was deprotected with 20% piperidine in dimethylformamide. The resin was cleaved with 95% trifluoroacetic acid, 2.5% triisopropylsilane and 2.5% dichloromethane (1 mL) for one hour. After ether precipitation, peptides were purified using reverse-phase HPLC. Final pure peptides were identified (≥95% purity) using LC/MS (Agilent Technologies 6120 Quadrupole LC/MS).

### Alignment analysis

Maximum Common Substructure (MCS) was carried out for alignment. The atom-atom pairing was obtained from the 2D maximum common substructure while the unpaired atoms are typed using extended atom types to enable relevant atomic overly. MCS and subsequent alignment was carried out using ChemAxon Similarity plugin.

### Membrane preparation

CHO-OPRD cells stably expressing β-arrestin 2 and δOR (DiscoverX, Fremont, CA) were grown in T75 flasks in 10ml F12 media (Fisher Scientific) containing 10% FBS (Sigma, Lot 12M246), 300μg/ml hygromycin B (Fisher Scientific) and 800μg/ml geneticin (Fisher Scientific) until confluency. Cells were dislodged by treatment with 0.25% trypsin/EDTA (Fisher Scientific) for 5 minutes. The trypsin was then deactivated with antibiotic-free F12 and the resulting cell suspension centrifuged for 5 minutes at 400G at room temperature (Eppendorf 5804R). The supernatant was aspirated and the pellet resuspended in 50 mM Tris HCl pH 7.4 (prepared from powder stock, Sigma) and sonicated on ice for 30 seconds (Qsonica XL-2000, level 3). The membranes were then isolated by ultracentrifugation for 30 minutes at 20,000 rpm (46,500g) in a SW41Ti rotor (Beckman Coulter Life Sciences, Indianapolis, IN) in a precooled Optima L-100 XP centrifuge (Beckman Coulter) at 4°C. The pellet was resuspended in 50 mM Tris HCl buffer and sonicated once more to homogenize. The suspension was then pulled through a 28G needle, aliquoted and frozen at −80°C for future use.

### Radioligand binding assay

For the binding assay 50 μl of a dilution series of peptide was added to 50 μl of 3.3 nM [^3^H]DPDPE (Kd =3.87 nM) in a clear 96 well plate. Next, 100 μl of membrane suspension containing 7 μg protein was added to the agonists and incubated for 90 minutes at room temperature. The reaction mixture was then filtered over a GF-B filter plate (Perkin Elmer) followed by 4 quick washes with ice-cold 50 mM Tris HCl. The plate was dried overnight after which 50 μl scintillation fluid (Ultimagold uLLT) was added and radioactivity was counted on a Packard TopCount NXT scintillation counter. All working solutions were prepared in a radioligand assay buffer containing 50mM Tris HCl, 10mM MgCl_2_, and 1mM EDTA at pH 7.4.

### Cell culture and biased signaling assays

cAMP inhibition and β-arrestin-2 recruitment assays were performed as previously described (Chiang et al., 2016). In brief, for cAMP inhibition assays HEK 293 (Life Technologies, Grand Island, NY, USA) cells (20,000 cells/well, 7.5 μl) transiently expressing FLAG-mouseδ-OR (van Rijn and Whistler, 2009; Whistler et al., 2001) and pGloSensor22F-cAMP plasmids (Promega, Madison, WI, USA) were incubated with Glosensor reagent (Promega, 7.5 μl, 2% final concentration) for 90 minutes at room temperature. Cells were stimulated with 5 μl δ-OR agonist 20 minutes prior to 30 μM forskolin (5 μl) stimulation for an additional 15 minutes, both occurring at room temperature. For β-arrestin recruitment assays, U2OS-humanδ-OR PathHunter β-arrestin-1 cells (DiscoverX) and CHO-humanδ-OR PathHunter β-arrestin-2 cells were plated (2500 cells/well, 10 μl) prior to stimulation with 2.5 μl δ-OR agonists for 90 minutes at 37°C/5%CO_2_, after which cells were incubated with 6 μl cell PathHunter assay buffer for 60 minutes at room temperature as per the manufacturer’s protocol. Luminescence for each of these assays was measured using a FlexStation3 plate reader (Molecular Devices, Sunnyvale, CA, USA).

### Statistical analysis

Each data point for binding and arrestin recruitment was run in duplicate, and for the cAMP assay in triplicate. For figure 2, the averages of each run were combined to provide a composite figure in favor of a representative figure. All data are presented as means ± standard error of the mean and were performed using GraphPad Prism 7 (GraphPad Software, La Jolla, CA). In order to calculate Log (τ/KA), we followed the operational model equation in Prism 7 as previously described (van der Westhuizen et al, 2014). Subsequently bias factors were calculated using leu-enkephalin as the reference compound.

## Results

### Synthesis of rubiscolin peptides

The rubiscolin, deltorphin II and leu-enkephalin peptides can all be found in nature (**Figure 1A**), however to ensure purity of the rubsicolin peptides, we elected to synthesize rather than extract these peptides [see methods for details]. HPLC-MS analysis of the produced rubiscolin 5 and 6 peptides revealed single, sharp elution peaks with >99% purity (RT = 5.68 and 6.19 minutes, respectively) displaying mass spectra with parent peaks characteristic of the [M+H]^+^ ions of our desired products (m/z = 620.4 and 767.4 for rubiscolins 5 and 6, respectively), enabling their use in cellular assays to characterize their pharmacology (**Figure 1B**).

**Figure 1.**
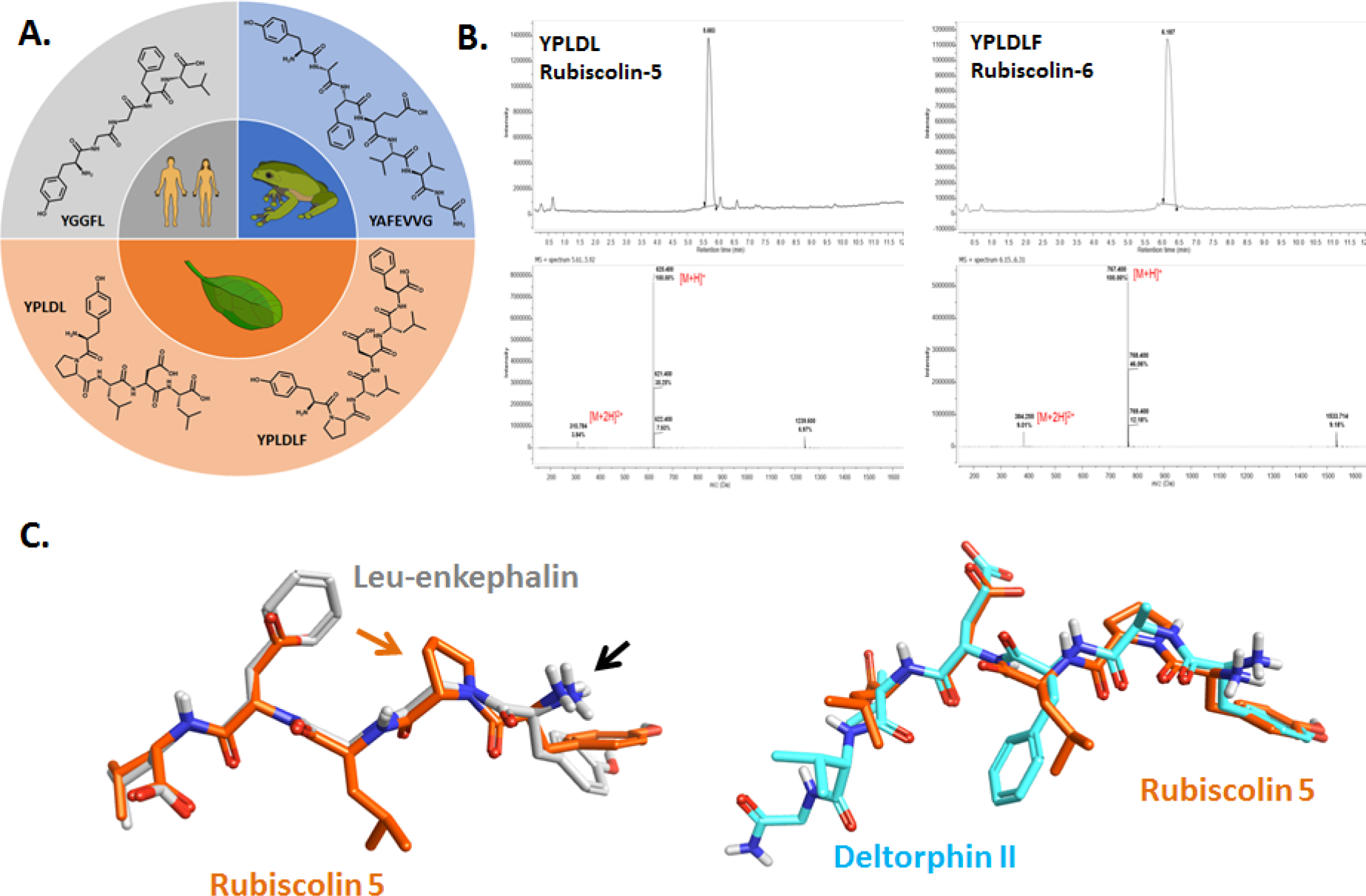
Origin and structural differences of naturally-occurring δ-OR peptides. **A**) Chemical structures of the naturally-occurring peptides leu-enkephalin (human), [D-Ala^2^]-deltorphin II (frog), rubiscolin-5 and rubiscolin-6 (spinach). **B**) HPLC-MS spectra for rubiscolin-5 and rubiscolin-6. **C**) Maximum common structure alignment for rubiscolin-5 (orange) with leu-enkephalin (gray) or deltorphin II (cyan). Black arrow points to the conserved amino-group. Orange arrow points to the proline residue unique to the rubiscolins.

### Maximal common substructure alignment of naturally occurring δ-OR peptides

The peptide ligands rubiscolin-5, and deltorphin II were subject to alignment using a maximal common substructure (MCS) method with leu-enkephalin. The conformation of leu-enkephalin used was derived from its crystal structure in complex with DPP3 (RCSB PDB ID: 5E3A). The MCS alignment between rubiscolin-5 and leu-enkephalin or deltorphin II provided similarity scores of 0.72 and 0.6 respectively, suggesting substantial overlap. Notable differences in the leu-enkephalin aligned rubiscolin-5 structure are the presence of an acidic aspartate in place of the hydrophobic phenylalanine of leu-enkephalin, and the presence of a bulky, hydrophobic leucine in place of leu-enkephalin’s glycine (**Figure 1C**). Interestingly, deltorphin II shows similar features to the rubiscolin compounds in this area with an acidic glutamate and hydrophobic phenylalanine (**Figure 1C**).

### Rubiscolin peptides are G-protein-biased δ-OR agonists

In order to determine the signaling properties of rubiscolin peptides at the δ-OR, we first confirmed that rubiscolin peptides indeed bind to the δ-OR with radioligand binding using membranes prepared from the PathHunter cells. We compared the affinity of the rubiscolin peptides to the highly potent and selective δ-OR peptides deltorphin II and leu-enkephalin. Deltorphin II displayed the strongest affinity for the δ-OR followed by leu-enkephalin, rubiscolin-6 and rubiscolin-5 (**Figure 2A**, **Table 1**).

**Table 1:** Pharmacological characterization of naturally occurring δ-OR peptides. Affinity of peptide agonists for the δ-OR is depicted as the pKi, which is a negative log value of the Ki, which is the concentration at which 50% of the δ-OR are occupied by the peptide. Potency and efficacy (α, normalized to leu-enkephalin) for the δ-OR peptide agonists to inhibit cAMP production is depicted as concentration of 50% inhibition (pIC_50_) and the SEM. Potency (pEC_50_) and efficacy (α, normalized to leu-enkephalin) of δ-OR agonists to recruit β-arrestin 1 and 2 are depicted with SEM. The number of repetitions for each drug is indicated in parentheses.

| Compounds | Affinity | cAMP | | β-arrestin1 | | β-arrestin2 | |
| --- | --- | --- | --- | --- | --- | --- | --- |
| | pKi | pIC<sub>50</sub> | $\alpha$ | pEC<sub>50</sub> | $\alpha$ | pEC<sub>50</sub> | $\alpha$ |
| **Leu-enkephalin** | 8.8±0.1 (4) | 9.0±0.1 (9) | 100 | 7.5±0.1 (11) | 100 | 7.5±0.1 (10) | 100 |
| **Deltorphin II** | 9.5±0.3 (5) | 9.4±0.1 (4) | 95±2 | 8.3±0.1 (5) | 94±6 | 8.7±0.2 (6) | 103±4 |
| **Rubiscolin-5** | 5.5±0.1 (4) | 5.2±0.1 (6) | 85±9 | ND (9) | ND | 4.1±0.3 (7) | 15±4 |
| **Rubiscolin-6** | 6.0±0.1 (3) | 5.8±0.2 (4) | 77±8 | ND (7) | ND | 5.1±0.3 (6) | 20±3 |

**Figure 2.**
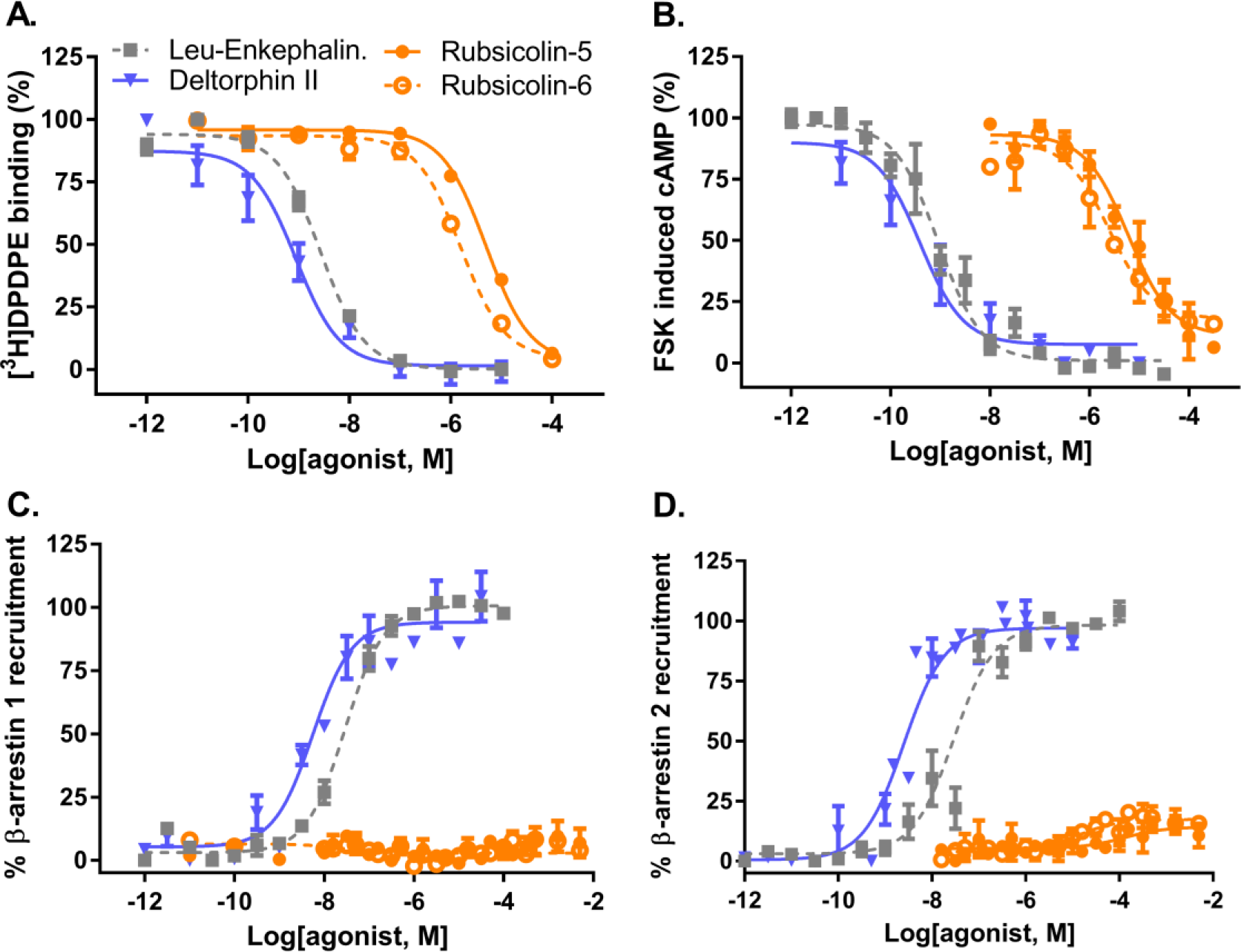
Pharmacological characterization of rubiscolin peptides at δ-ORs. **A**) Displacement of [^3^H]DPDPE from δ-ORs (inset shows saturation binding for [^3^H]DPDPE on δ-ORs). **B**) Inhibition of forskolin-induced cAMP production in HEK293-δ-OR cells. **C**) β-arrestin 1 recruitment in U2OS δ-OR PathHunter cells. **D**) β-arrestin 2 recruitment in CHO δ-OR PathHunter cells. Composite figures are shown (see Table 1 for details).

Next we confirmed that all four peptides were indeed agonists able to inhibit forskolin-induced cAMP production by activation of the inhibitory Gαi-protein pathway. All four peptides displayed full agonism and followed the same rank order in potency as seen for affinity (**Figure 2B**, **Table 1**). Next, we assessed the ability of these peptides to recruit β-arrestin, finding that deltorphin II and leu-enkephalin recruit both β-arrestin 1 (**Figure 2C**) and β-arrestin 2 (**Figure 2D**), with deltorphin II showing a higher potency (**Table 1**). Interestingly, in both assays β-arrestin recruitment induced by the rubiscolins ranged from low to undetectable even at concentrations of 316 μM (**Figure 2C and 2D**). Importantly, because the PathHunter cell line was used for our binding studies, this limited β-arrestin recruitment cannot be attributed to a lack of binding to the δ-OR.

## Discussion

Delta opioid receptor peptides like leu-enkephalin, DPDPE and DADLE are all largely unbiased; still, G-protein biased synthetic δ-OR peptides have been produced over 10 years ago as exemplified by UFP-512 (Chiang et al., 2016; Molinari et al., 2010). We find that deltorphin II has a bias towards β-Arrestin recruitment (**Table 2**, bias factor < 1). Distinguishingly, here we present the first report of rubiscolin-5 as naturally-occurring G-protein-biased δ-OR peptide, with a bias factor 2 when comparing cAMP versus β-arrestin 2 (**Table 2**) and undetectable rubiscolin-5 induced β-arrestin 1 recruitment to the δ-OR at concentrations as high as 2mM (**Figure 2C and D**). Rubiscolin-6 similarly did not recruit β-arrestin 1 to an extent that would allow us to calculate a bias factor. However the cAMP-β-arrestin 2 bias factor for rubiscolin-6 was <1 suggestive of β-Arrestin-bias. This appears surprising given that at a 10 μM concentration 80% of δ-ORs will have bound rubiscolin-6 causing 66% inhibition of cAMP yet only 5% β-arrestin 2 recruitment (**Figure 2**). It should be noted that potency, and particularly potency difference between the test compound and reference compound carries a lot of weight relative to efficacy when calculating bias factor (Brust et al., 2015).

**Table 2:** Bias factors for naturally occurring δ-OR peptides. Transduction coefficient (t), (K_A_) and bias factor determination. Log (τ/K_A_) was determined using the operational model equation (van der Westhuizen et al, 2014). ΔLog (τ/K_A_) was calculated using Leu-enkephalin as the reference compound. Bias Factor = 10^(ΔLog (τ/KA)[cAMP]−ΔLog (τ/KA)[βARR])^.

| Compounds | cAMP | | β-arrestin1 | | β-arrestin2 | | Bias Factor | |
| --- | --- | --- | --- | --- | --- | --- | --- | --- |
| | Log ($\tau/K_A$) | $\Delta\text{Log}(\tau/K_A)$ | Log ($\tau/K_A$) | $\Delta\text{Log}(\tau/K_A)$ | Log ($\tau/K_A$) | $\Delta\text{Log}(\tau/K_A)$ | cAMP-βarr 1 | cAMP-βarr 2 |
| Leu-enkephalin | 9.05 | 0 | 7.56 | 0 | 7.53 | 0 | 1 | 1 |
| Deltorphan II | 9.43 | 0.39 | 8.20 | 0.64 | 8.56 | 1.07 | 0.55 | 0.21 |
| Rubiscolin-5 | 5.13 | -3.92 | ND | NA | 3.26 | -4.23 | NA | 2.04 |
| Rubiscolin-6 | 5.41 | -3.63 | ND | NA | 4.14 | -3.35 | NA | 0.52 |

ND = not determinable; NA = not applicable

In terms of opioid receptor pharmacology, studies thus far have highlighted the benefits of biased signaling that favors G-protein over β-arrestin. For μ-ORs, G-protein biased signaling may reduce tolerance, constipation and respiratory depression (DeWire et al., 2013; Manglik et al., 2016), although recent studies have placed some doubts about the strength of this hypothesis (Altarifi et al., 2017; Hill et al., 2018). For the κ-ORs, G-protein biased ligands may avoid aversive effects commonly associated with (unbiased) κ-OR agonists (Bruchas et al., 2007; Land et al., 2009). For the δ-ORs β-arrestin recruitment has been associated with increased alcohol use, whereas G-protein-biased δ-OR agonists reduce alcohol intake in mice (Chiang et al., 2016; Robins et al., 2018).

Currently only an antagonist-bound δ-OR structure has been resolved, in contrast to κ-ORs and μ-ORs for which agonist-bound structures are available. Still, although less optimal, homology modeling can be performed to use the agonist-bound OR structures to create a computational model of an agonist bound conformation of the δ-OR. Recent studies using computational modeling and X-ray crystallography have suggested that drug conformation can have a significant impact on specific receptor conformations inducing receptor states that are more or less susceptible to recruit β-arrestin (McCorvy et al., 2018; Wacker et al., 2017). While common substructure analysis did not show major differences in conformation between rubiscolin-5 and leu-enkephalin or deltorphin II, it is noteworthy that the rubiscolin peptides contain a proline residue. Proline residues, due to their cyclic structure are known to induce kinks and instill a degree of inflexibility to the peptide. It is possible that this proline residue in rubiscolins induces a certain conformation that the more flexible enkephalin and deltorphin II circumvent. With our finding that rubiscolin peptides are G-protein-selective δ-OR agonists, it may be of interest to dock rubiscolin-6, or YPLDLV a more potent and δ-OR-selective synthetic analog (Yang et al., 2003b) to potentially obtain insight into which regions of the δ-OR are being engaged or spared to confer G-protein selectivity onto rubiscolin when compared with deltorphin II or leu-enkephalin.

Rubsicolin-6 is more potent and has stronger affinity for the δ-OR than rubiscolin-5, but both peptides are rather weak in comparison to leu-enkephalin and deltorphin II, which may limit their therapeutic potential. One advantage however of the rubisocolin peptides is their oral bioavailability; rubiscolins are produced by digestive pepsin cleavage of rubisco, and rubiscolin-6 at a dose of 100 mg/kg, p.o. produced analgesia in a tail-pinch assay (Yang et al., 2001), enhanced memory consolidation, (Yang et al., 2003a) and reduced anxiety-like behavior (Hirata et al., 2007) in a manner that was shown to be reversible by the δ-OR antagonist naltrindole, suggesting that the behavior was δ-OR mediated.

Rubiscolin-6 has been reported to produce an orexigenic response (Kaneko et al., 2012a; Kaneko et al., 2014; Kaneko et al., 2012b) by acting at δ-ORs, however DPDPE, a peptide which recruits β-arrestin on par with enkephalin, produces a similar physiological response (Kaneko et al., 2014), suggesting that the G-protein-selective nature of rubiscolin-6 may not be a contributing factor to this physiological response.

Overall, we provide evidence for the first known naturally-occurring, orally bioavailable G-protein-selective δ-OR peptide agonist. The attributes of rubiscolin-6 may make it an interesting drug in treating alcohol use disorder (Chiang et al., 2016; Robins et al., 2018), or as an analgesic with fewer side effects.

## Author contributions

RJC and KLM performed the cellular pharmacological assays and BZ synthesized and ran quality control for the rubiscolin peptides. AHM ran the peptide alignments. RJC, KLM and RMvR. designed the study, and drafted the manuscript. RJC, KLM and RMvR analyzed the data. RMvR supervised RJC and KLM. DT supervised BZ. MAL supervised AHM. All authors proofread and approved the final draft of the manuscript.

